# Biofilm Growth Under Elastic Confinement

**DOI:** 10.1101/2021.12.05.471265

**Authors:** George T. Fortune, Nuno M. Oliveira, Raymond E. Goldstein

## Abstract

Bacteria often form surface-bound communities, embedded in a self-produced extracellular matrix, called biofilms. Quantitative studies of their growth have typically focused on unconfined expansion above solid or semi-solid surfaces, leading to exponential radial growth. This geometry does not accurately reflect the natural or biomedical contexts in which biofilms grow in confined spaces. Here we consider one of the simplest confined geometries: a biofilm growing laterally in the space between a solid surface and an overlying elastic sheet. A poroelastic framework is utilised to derive the radial growth rate of the biofilm; it reveals an additional self-similar expansion regime, governed by the stiffness of the matrix, leading to a finite maximum radius, consistent with our experimental observations of growing *Bacillus subtilis* biofilms confined by PDMS.

---

Bacterial biofilms are microbial accretions, enclosed in a self-produced polymeric extracellular matrix [1], which adhere to inert or living surfaces. A biofilm gives the individual cells a range of competitive advantages, such as increased resistance to chemical attack. Since the popularisation in the mid 1600s of the light microscope as a tool to study problems in biology [2, 3], observations of groups of bacteria on surfaces have been amply documented [4], most notably by van Leeuwenhoek in his dental plaque [5]. Yet, it is only in the last few decades with the development of new genetic and molecular techniques that the complexity of these communities has been appreciated and biofilm formation has been recognised as a regulated developmental process in its own right [6, 7].

Biofilm formation is common across a wide range of organisms in the archaeal and bacterial domains of life, on almost all types of surfaces [8]. Cells attach to a surface and form micro-colonies through clonal growth. These then grow and colonise their surroundings through twitching motility [1]. A central research focus has been understanding these growth dynamics. Building on important work on osmotically-driven spreading [1], a biofilm has often been modelled as a viscous, Newtonian fluid mixture (nutrient rich water and biomass), neglecting the matrix elasticity. The effects of surface tension [10], osmotic pressure [11], and the interplay between nutrients, cell growth, and electrical signaling in response to metabolic stress have all been studied recently [2].

While previous analyses have focused on the experimentally tractable cases of unconfined and unsubmerged biofilms [1, 2, 10, 11], they do not accurately reflect the conditions in which many biofilms grow; they thrive in confined micro-spaces [13] between flexible elastic boundaries such as vessel walls or soil pores [14], and indeed in the human body, where they account for over 80% of microbial infections [15]. Biofilms are difficult to treat with antibiotics, being thousands of times more resistant than the constituent microorganisms in isolation [16] due to a range of mechanical and biological processes [17, 18]. The recent rapid growth in the use of implantable biomedical devices (stents, catheters, and cardiac implants) has brought with it a large increase in associated biofilm infections [19] since artificial surfaces require much smaller bacterial loads for colonisation than the corresponding volume of native tissue (≈ 10^−4^ as much [20]).

Here we develop the simplest model for a confined biofilm, using a poroelastic framework to obtain a system of equations describing its expansion dynamics. We find an analytic similarity solution for the biofilm height and radius, together with the vertically averaged biomass volume fraction. Consistent with experimental observations on growing *Bacillus subtilis* biofilms described here, unlike unconfined biofilms whose radius grows exponentially, the balance between elastic stresses and osmotic pressure difference across the interface implies an additional possible growth regime where within a shallow layer lubrication assumption, confined biofilms have a maximum radius at long times. The transition between regimes is governed by the stiffness of the matrix.

We consider a bio-mechanical system in which bacteria grow and divide, converting nutrient-rich fluid into biomass and thus inducing a flow of biomass outwards from the biofilm centre. This flow is resisted by elastic stresses within the extracellular matrix (ECM), while the biofilm height dynamically adjusts to ensure conservation of normal stress across the overlying elastic sheet. An influx of water assures volume conservation. Illustrated in Fig. 1, an axisymmetric biofilm of thickness *h*(*r, t*), radius *R*(*t*) and biomass volume *V* rests on an impermeable flat plate at *z* = 0 and grows below an elastic sheet of thickness *d* = 𝒪(*R*) and bending modulus *B* = *Ed*^3^*/*12(1 − *ν*^2^), where *E* and *ν* are the Young’s modulus and Poisson’s ratio of the sheet. We examine the simplest biofilm composition, a mixture of bacteria (volume fraction *ϕ*_*b*_), sugar-rich secreted polymeric ECM (volume fraction *ϕ*_*m*_), and nutrient-rich water (modelled as a low viscosity Newtonian fluid [1] with dynamic viscosity *μ*_*f*_ and volume fraction 1 − (*ϕ*_*m*_ + *ϕ*_*b*_) ≡ 1 − *ϕ*), under the assumption that *ϕ*_*m*_ ≪ *ϕ*_*b*_ [1]. For theoretical simplicity, we assume that the biomass volume fraction *ϕ* is independent of *z*, so *∂ϕ/∂z* = 0.

**FIG. 1.**
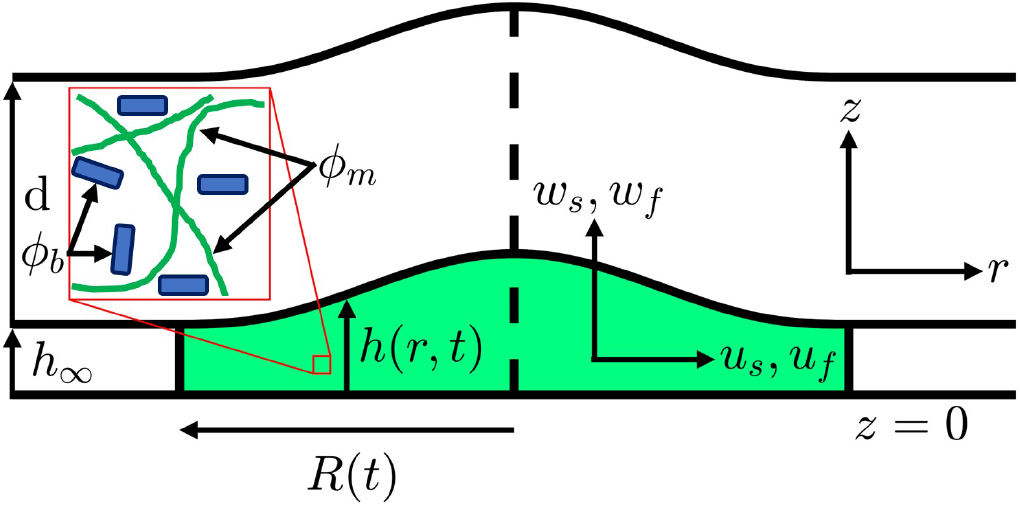
Schematic of a confined biofilm. An axisymmetric biofilm (green) grows between a rigid surface at *z* = 0 and an elastic sheet at *z* = *h*, with undeformed gap height *h*_∞_. Inset: the biomass is a mixture of bacterial cells (blue, volume fraction *ϕ*_*b*_) and extracellular matrix (green,volume fraction *ϕ*_*m*_). The pore-averaged velocities of the solid and fluid phases are denoted by ***u***_***s***_ = (*u*_*s*_, *w*_*s*_) and ***u***_***f***_ = (*u*_*f*_, *w*_*f*_).

We denote the pore-averaged velocity and stress tensor of the solid and liquid phases by {***u***_***s***_ = (*u*_*s*_, *w*_*s*_), ***σ***_***s***_} and {***u***_***f***_ = (*u*_*f*_, *w*_*f*_), ***σ***_***f***_ ≈ −*p****I***} [1] respectively, where *p*, Π and 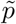 are the pore, osmotic, and bulk pressures (with 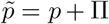[21]). Since the vertical deflection of the sheet Δ*d* =𝒪(*h*) is small compared to its thickness *d*, we ignore stretching and model it as a thin elastic beam with radius of curvature 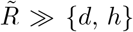 and surface tension *γ* against the biofilm. We neglect gravity, assume that nutrient concentrations across the biofilm are constant, and take the biomass growth rate *g* to have the saturating form

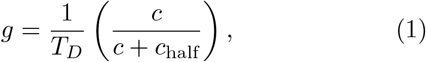

independent of position, where *T*_*D*_ is the doubling time (typically hours), *c* is the concentration of a limiting nutrient and *c*_half_ is that for half-maximum growth rate. Both *c* and hence *g* are taken to be constant in light of our experiments, introduced below, in which there is an external flow that ensures homogeneity. Conserving mass in both the solid and fluid phases gives

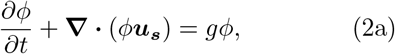

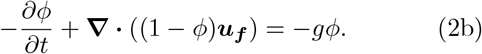

Defining the Terzaghi effective stress tensor as ***σ*** = *ϕ*(***σ***_***s***_ − ***σ***_***f***_) [22], momentum balance yields

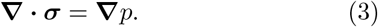

To model ***σ***, we deviate from prior work that assumed a Newtonian fluid by adopting a poroelastic framework that incorporates the elasticity of the ECM. In this picture, ***σ*** obeys the elastic constitutive law

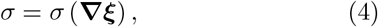

where ***ξ*** = (*ξ, ζ*), the deformation vector of the medium away from a reference state, is related to the biofilm phase velocity through ***u***_***s***_ = (*∂*_*t*_ + ***u***_***s***_ **· ∇**) ***ξ***. Little utilised in the study of biofilms, it is a common approach in many problems containing elasticity in geophysics (hydrology subsidence and pumping problems [23, 24] or industrial filtration [25]) and biological physics (cell cytoplasm [26]). Here, we consider the simplest case, where ***σ*** obeys the linear constitutive law

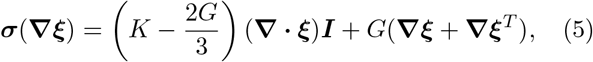

where *K* and *G* are the effective bulk and shear moduli of the biofilm respectively, assumed constant. As in [23], *K* and *G* are properties of the whole biofilm rather than just the ECM. We prescribe explicitly the general form for the horizontal velocity of the solid phase,

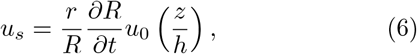

where *u*_0_ is the *z*−dependent part of *u*_*s*_. We take

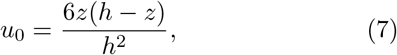

since this is the simplest functional form obeying no-slip boundary conditions at *z* = 0 and *z* = *h* as well as ⟨*u*_0_⟩ = 1. However, as shown below, we find a solution independent of the exact form for *u*_0_. Global volume conservation gives *∂R/∂t* while *r/R* sets a simple linear radial dependence, ensuring that *u*_*s*_ = 0 at *r* = 0. As for *u*_0_, tweaking this radial dependence does not qualitatively change the resulting dynamics of the system.

In contrast, vertical flow is governed by pressure gradients induced both by the upper confinement and by elastic stresses in the extracellular matrix. We invoke Darcy’s law for flow within the matrix, giving

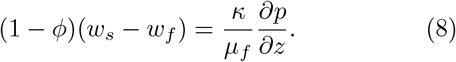

where *κ* = *κ*(*ϕ*) is the effective biofilm permeability with characteristic permeability scale *κ*_0_. The osmotic pressure away from equilibrium Π(*ϕ*) is taken to be that of-Flory Huggins theory [27], with interaction parameter *χ* ≃ 1*/*2 so there is no demixing [28]. Assuming that the matrix solid fraction *β* = *ϕ*_*m*_*/ϕ* ≪ 1 is constant across the biofilm, the osmotic pressure is [29]

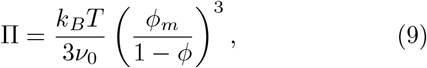

a function of thermal energy *k*_*B*_*T* and *ν*_0_, the effective volume occupied by one monomer of matrix. Since the matrix consists of many different substances, notably sugars, proteins and DNA, we estimate *ν*_0_ by the volume occupied by one sugar monomer. This term is subdominant in the analysis below, and thus does not appear in the interior (*r* ≤ *R*) solutions (14) - (18). We close this system of equations with a set of vertical boundary conditions, given in the Supplementary Material [30].

The analysis exploits two separations of scales: (i) the initial radius of the confined biofilm *R*_0_ = *R*(*t* = 0) is much greater than the initial height *H*_0_ = *h*(*r* = 0, *t* = 0), a lubrication approximation, and (ii) the growth time scale 1*/g* is much larger than the poroelastic equilibration time 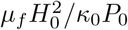. We nondimensionalise the equations anisotropically using these length scales, denote the vertically averaged form of a function *f* by 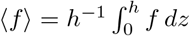, and define *φ* = ⟨ *ϕ⟨, υ*_*s*_ = ⟨*u*_*s*_⟩, *k* = ⟨*κ*⟩, 𝒫 = *p/P*_0_ and

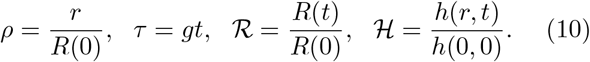

Keeping only leading-order terms in *ϵ* = *H*_0_*/R*_0_ [30], the model reduces to coupled PDEs for the height ℋ(*ρ, τ*) and depth-averaged biomass fraction *φ*(*ρ, τ*) as functions of radial distance *ρ* and time *τ*. The horizontal pressure gradient adjusts to one of three possible modes

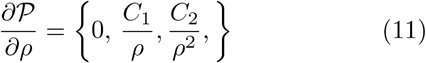

where *C*_1_ and *C*_2_ are constants and the dominant contribution to the pressure 𝒫 arises from the bending stresses imposed from the upper elastic sheet,

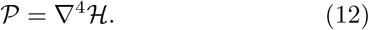

The depth-integrated biomass fraction *φ*ℋ satisfies a conservation law of the form *∂*(*φ*ℋ)*/∂τ* = −**∇** · **𝒥**_*φ*_ + **𝒮**,

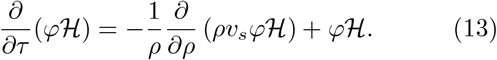

Thus, *φ*ℋ grows exponentially from the source term **𝒮** = *φ*ℋ, while subject to radial advection at speed *υ*_*s*_(ℋ,*ℛ*) from the flux term **𝒥**_*φ*_. The system is closed with a set of boundary conditions, deriving the boundary conditions for ℋ at the biofilm interface by extending the framework outside the biofilm to the whole domain and imposing far field boundary conditions [30]. In the mode zero case when the horizontal pressure gradient is zero, Eqs. (11)-(13) admit the interior (*ρ* ≤ ℛ) solutions

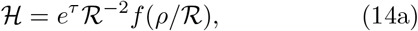

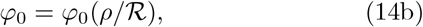

where

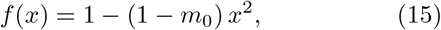

the incline ratio

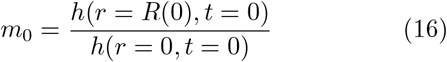

is a measure of the initial flatness of the biofilm, *φ*_0_(*ρ*) = *φ*(*ρ, τ* = 0) is set from the initial conditions and we have utilized the vertically-averaged boundary conditions [30] and the initial conditions ℋ(*ρ* = 0, *τ* = 0) = ℛ (*τ* = 0) = 1 and ℋ(*ρ* = 1, *τ* = 0) = *m*_0_. The form of (14) guarantees that the total biomass *∫dρρ* ℋ*φ* grows as e^*τ*^. We obtain ℛ (*τ*) as the solution of the cubic equation

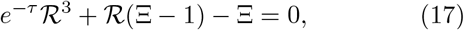

where the single free parameter is

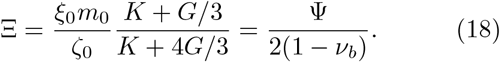

Derived in [30], Ψ = *ξ*_0_*m*_0_*/ζ*_0_ is a measure of the initial ratio between horizontal and vertical stress gradients in the biofilm while *ν*_*b*_, the effective Poisson’s ratio of the ECM, is a measure of how stiff the biofilm is (stiffer biofilms have lower *ν*_*b*_). The radial expansion of the biofilm is mediated by a balance at the biofilm edge between horizontal and vertical elastic deformation in the biofilm (the Ξ and *e*^−*τ*^ℛ^3^ terms, respectively, in (17)) and the osmotic pressure difference across the biofilm interface (the ℛ (Ξ − 1) term).

For general Ξ and *τ*, this equation does not always admit an analytic solution and is solved numerically [30]. Figure 2(a) plots the temporal evolution of ℛ for a range of different values of Ξ. Figure 2(b) explores this further, choosing a fixed observation time *τ*_0_ and plotting ℛ(*τ*_0_) as a function of Ξ. Two clear regimes emerge. If Ξ *<* 1, the first and second terms in (17) dominate in a balance between stresses caused by the vertical elastic deformations and the osmotic pressure difference, leading to a limit on vertical expansion. The biofilm then spreads with exponential radial growth [1], with ℛ → (1 − Ξ)^1*/*2^ *e*^*τ/*2^ as *τ* → ∞. If Ξ *>* 1 (the dark blue curves in figure 2(a)), the second and third term in (17) are dominant, giving a balance between stresses caused by horizontal elastic deformations and the osmotic pressure difference that limits horizontal expansion. The radius at intermediate times exhibits power-law growth before slowing down to reach a maximum ℛ(∞) = Ξ*/*(Ξ − 1), when the shallow layer approximation is still valid. In the special case Ξ = 1, the osmotic pressure difference across the interface is zero, leading to a balance between horizontal and vertical elastic stresses. As shown in Fig. 2(a), the system exhibits transitional exponential growth, with ℛ = *e*^*τ/*3^, but this state is not stable; curves with Ξ just above and below unity will veer off eventually to tend to a constant radius or to the faster *e*^*τ/*2^ growth law.

**FIG. 2.**
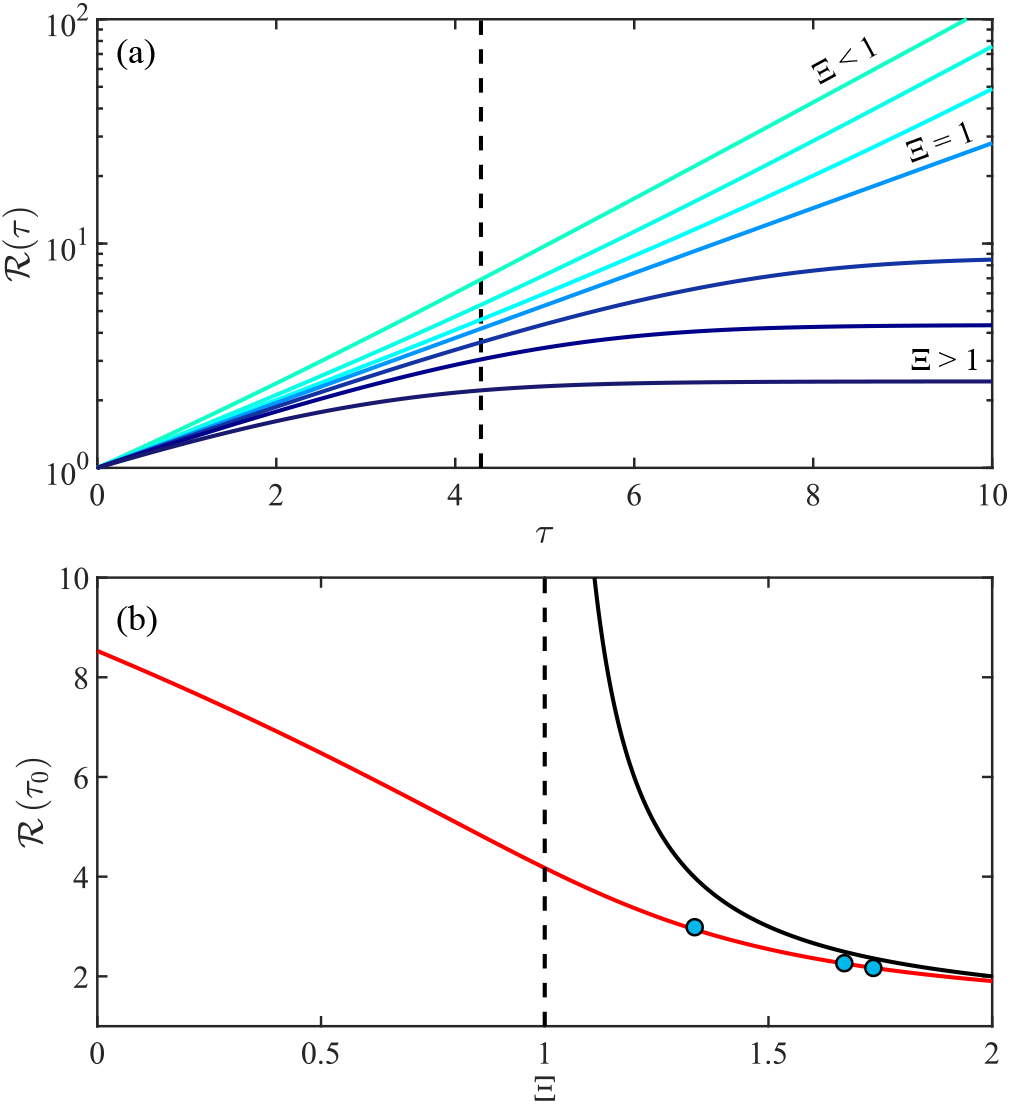
Growth dynamics of confined biofilms according to the poroelastic model. (a) The scaled biofilm radius ℛ as a function of scaled time in a semilogarithmic plot, for Ξ ∈ [0.4, 0.75, 0.91, 1, 1.13, 1.3, 1.7]. Darker colours denote larger Ξ. (b) Biofilm radius at a fixed *τ*_0_ (dashed vertical line in (a)) as a function of Ξ, both numerically (**-**) and experimentally (•), and numerically for *τ*_0_ → ∞ (**-**).

We performed experiments on the growth of biofilms confined by polydimethylsiloxane (PDMS), the results of which can be compared directly to the model developed above. The methodology follows existing protocols [2–4] developed to understand the growth of focal (and submerged) biofilms under well-defined flow conditions. Full details are given in Supplemental Material [30]; here we summarize the key features. Flagella-less mutants of *Bacillus subtilis* were used to avoid secondary contributions to biofilm spreading [1]. Cells in exponential growth phase were centrifuged and resuspended in growth medium before being loaded at the centre of Y04-D plates linked to the CellASIC ONIX microfluidic platform (EMD Millipore), and kept at 30 ^*°*^C. In this setup, they are confined between glass and an overlying PDMS sheet of thickness *d* = 114 *μ*m, with an initial gap of *h* = 6 *μ*m. Fresh medium was flowed through the chamber with a mean speed of ∼ 16 *μ*ms^−1^ [2–4]. Biofilm growth was imaged at 1 frame/minute on a spinning-disc confocal microscope in bright field. As the biofilms were often frilly, with long thin strands of matrix polymer protruding from their edges, a Gaussian image processing filter in MATLAB was used to neglect these strands when identifying the interface with a Sobel edge detector.

Figure 3(a) is a montage of the expanding biofilm edge and the best-fit circle for one particular experiment, while Figure 3(b) plots the scaled biofilm radius ℛ as a function of time. In a clear departure from unconfined bacterial biofilms, the ℛ initially grows as a power law before tending to saturate at long times. These profiles exhibit the main qualitative features predicted by the theoretical model for Ξ *>* 1. The lines of best fit (black lines in 3(b), [30]) show good agreement over the entire time course of the experiments. A further comparison with theory is obtained by measuring in three different experiments, at the same nutrient concentration, the radius ℛ(*t*_0_) at a particular time *t*_0_ = 5 h, chosen as a time when the biofilm radius had a least doubled from its initial value. The parameter *g* relating absolute and rescaled times was fitted across all experiments, and gives the value *τ*_0_ = 4.29 used in Fig. 2(b), while Ξ is fitted independently for each. These experimental points in the Ξ − ℛ plane are shown as blue circles in Figure 2b), and agree very well with the poroelastic model developed here.

**FIG. 3.**
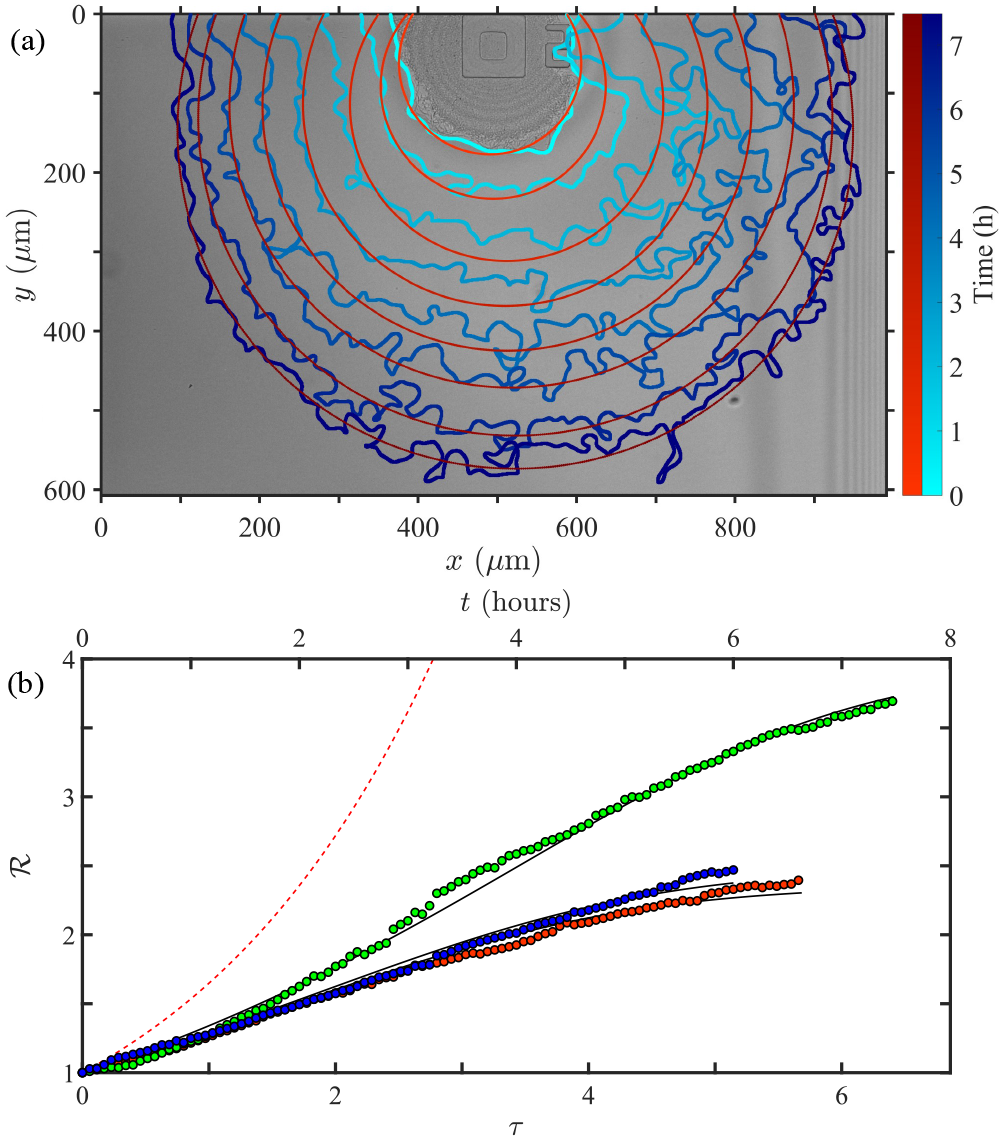
Experimental growth of *B. subtilis* biofilms under confinement by a PDMS sheet. (a) Montage plot, superimposed on image of the initial biofilm, showing the temporal evolution of the biofilm boundary (blue curves; darker colors denote later times) and fitted circles (red). (b) Scaled biofilm radius ℛ against scaled time for 3 experiments (•,•,•) compared to fitted dynamics from model (**–**) Dashed red curve is ℛ(*τ*) = e^*τ/*2^ expected for growth at constant thickness.

Motivated by the desire to understand the evolution of biofilms under confinement, we have constructed a minimal mathematical model that uses a poro-elastic framework. This admits a family of self-similar quasi-steady solutions, parameterized by a dimensionless parameter Ξ that measures the elasticity of the matrix. Those solutions are consistent with the experimentally observed behavior of confined *B. subtilis* biofilms. For comparison, [30] presents the corresponding theoretical model in which, following previous work in the literature, the biomass is modelled instead as a viscous Newtonian fluid, neglecting the intrinsic elasticity of the biofilm ECM. In that case, a solution with power law growth tending to a maximum finite biofilm radius is not supported, demonstrating that modelling the matrix elasticity is essential to capturing biofilm growth under elastic confinement.

Unlike unconfined biofilms, a subset of these solutions (where Ξ *>* 1) have a maximum radius due to a balance between elastic stresses and the osmotic pressure difference across the interface. The key parameter that determines which regime the system lies in and thus whether the biofilm grows predominately radially or axially is the stiffness of the biofilm matrix. Hence, we may view matrix elasticity is a competitive trait that could well be optimized by natural selection.

We are grateful to G.G. Peng and J.A. Neufeld for discussions and P.A. Haas and A. Chamolly for valuable comments on an earlier version of the manuscript. This work was supported in part by the Engineering and Physical Sciences Research Council, through a Doctoral Training Fellowship (GTF) and an Established Career Fellowship EP/M017982/1 (REG), and by a Wellcome Trust Interdisciplinary Fellowship and Discovery Fellowship BB/T009098/1 from the Biotechnology and Biological Sciences Research Council (NMO).

## Supplementary Material

We present dimensionless shallow layer scalings that reduce the system of equations describing the full system (2)-(6) in the main text to a pair of coupled differential equations for the height *h*(*r, t*) and vertically averaged biomass volume fraction ⟨*ϕ*⟩ (*r, t*) as a function of radial distance *r* and time *t*. The deformation ***ξ*** is expressed as a function of derivatives of *h*, utilizing both global biomass volume conservation and a pressure condition at the biofilm interface. The system is closed with boundary conditions for *h* at the biofilm interface, obtained by extending the framework outside the biofilm to the whole domain and imposing far-field free-beam and zero-pressure conditions.

### EXPERIMENTAL

#### Supplementary information on methods and materials

All experiments reported here used flagella-null cells of *Bacillus subtilis* (NCIB 3610 hag::tet, a gift from Roberto Kolter). Flagellaless cells were preferred because their inability to swim largely avoids contamination of inlets loaded with fresh growth medium in the microfluidic devices, and removes motility as a secondary contribution to biofilm spreading, as in earlier work [S1].

For each experiment, *Bacillus subtilis* cells were streaked from −80^°^C freezer stocks onto 1.5% agar LB plates and incubated at 37^°^C for 12 hours. Cells from a single colony were then inoculated in LB Broth (Lennox) at 37^°^C for 3 hours to obtain cells in the exponential growth phase. These were centrifuged at 2600 rpm for 6 minutes and re-suspended with fresh minimal salts glycerol glutamate (MSgg), the standard biofilm growth medium for *B. subtilis* [S1]. This MSgg medium contained 5 mM potassium phosphate buffer (pH 7.0), 100 mM MOPS buffer (pH 7.0), 2 mM MgCl_2_, 700*μ*M CaCl_2_, 50*μ*M MnCl_2_, 100 *μ*M FeCl_3_, 1 *μ*M ZnCl_2_, 2 *μ*M thiamine HCl, 0.5% (v/v) glycerol and 0.5% (w/v) monosodium glutamate.

Cells were then loaded at the center of Y04-D plates linked to the CellASIC ONIX microfluidic platform (EMD Millipore), and were incubated at 30 ^°^C for the duration of each experiment. In this setup, they were confined between a rigid surface (glass) and an elastic sheet (PDMS, 114 *μ*m thick), a distance 6 *μ*m apart. In all experiments we flowed fresh MSgg medium via one inlet, using a pump pressure of 1 psi, corresponding to a mean flow rate of 16 *μ*ms^−1^ in the growth chamber [S2–S4]. The subsequent growth of these submerged biofilms was then followed over time with a Zeiss Axio Observer Z1 microscope, connected to a Yokogawa Spinning Disk Confocal CSU and controlled by Zen Blue software. A Zeiss 10 × */*0.3 M27 Plan-Apochromat objective lens was used to acquire bright-field images at a rate of 1 frame per minute. These images were analyzed using both the open source image processing package Fiji [S5] and several custom MATLAB scripts utilizing MATLAB’s Image Processing Toolbox. In particular, a Sobel edge detector was used to locate the biofilm edge. The experimental biofilms were often frilly with long thin strands of matrix polymer protruding from the biofilm edge. Hence, 2D gaussian filtering using the MATLAB inbuilt function imgaussfilt was used to neglect these strands when identifying where the interface is. In order to fit a circle to the extracted interface a least-squares fit was implemented.

#### Raw experimental data

Figure S1 gives the corresponding raw data for the experiment given in the montage plot of Figure 3(a) of the main text, showing how in Figure S1(a) the scaled biofilm radius ℛ and in Figure S1(b) *σ*_*b*_, a measure of the circularity of the biofilm edge, vary with time, where *σ*_*b*_ satisfies

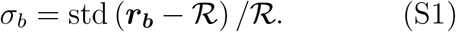

Here, ***r***_***b***_ is a vector giving the scaled distance of the points on the biofilm edge from the center of the biofilm. As a biofilm grows, it becomes more circular (after an initial increase due to growth around an obstacle *σ*_*b*_ decreases monotonically) but with frillier edges. Furthermore, as shown in the montage plot, interference fringes (Newton rings) are used to gain a qualitative understanding of how the upper PDMS sheet deforms. In particular, the fringes are circular, implying that the sheet deforms asymmetrically and thus evolves consistently with one of the key assumptions of the theoretical model, namely that *h* = *h*(*r, t*) is independent of *θ*.

**FIG. S1.**
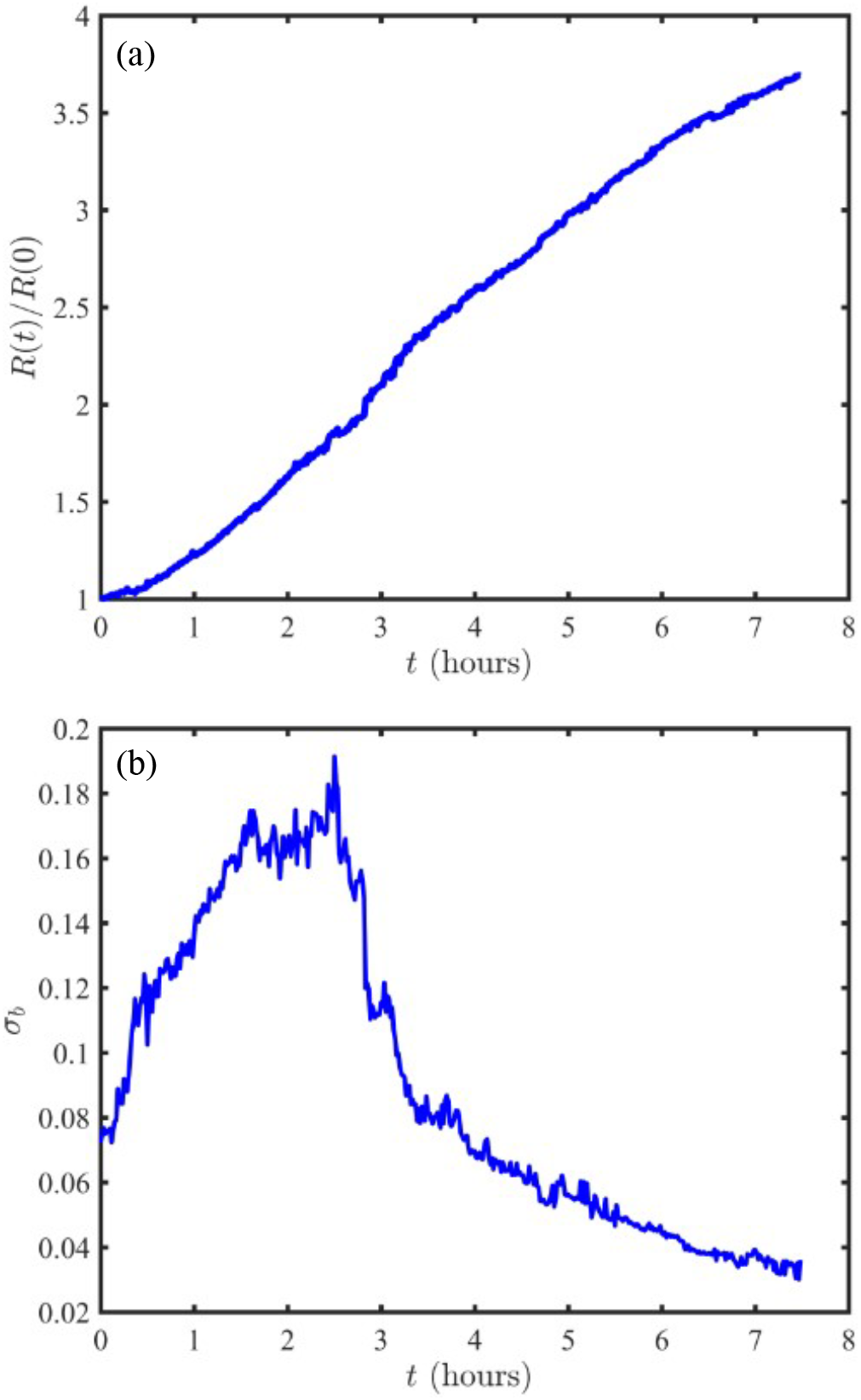
Raw data showing for a particular experiment how the scaled biofilm radius ℛ (a) and the relative deviation of the biofilm interface from a least-squares fitted circle *σ*_*b*_ (b) vary as functions of time.

#### Fitting Procedure

Numerical solutions of (17) predict the evolution of ℛ as a function of dimensionless time *τ*, with a single fitted parameter Ξ. To convert back to real time, the biofilm growth timescale *τ*_0_ = *g*^−1^ has to be determined. This was found through an iterative procedure, utilising all three experimental datasets to obtain a series of increasingly accurate estimates for *g*, {*g*_1_, *g*_2_, …}, using the recursion relation that *g*_*n*+1_ is the *g* that minimises

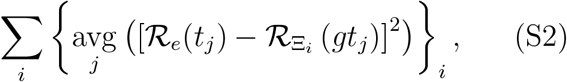

where *i* iterates over all datasets and *j* over all points within the experimental dataset ℛ_*e*_(*t*_*j*_) = *R*_*e*_(*t*_*j*_)*/R*_*e*_(0) enumerated by 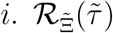 is the solution to (17) that is numerically computed for 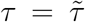 and 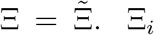 is the value of Ξ that for the data set enumerated by *i* minimises the objective function

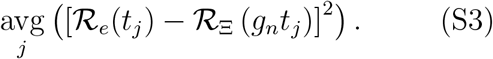

Here, all minimizations were performed using the MATLAB inbuilt function fminbnd [S6]. This resulted in fitted values for the biofilm growth time scale of *g* = 0.8574 and for Ξ of 1.7352, 1.6702 and 1.3358 for the three different experiments.

### FULL POROELASTIC FRAMEWORK

Below, we denote the region which the biofilm occupies (*r* ≤ *R*) the inner region and the region outside of the biofilm (*r* ≥ *R*) the outer region.

#### Inner Dimensional Vertical Boundary Conditions

Since horizontal motion of the upper PDMS sheet can be neglected, imposing no-slip boundary conditions at both the lower and upper boundaries yields

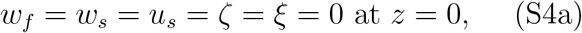

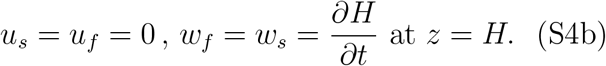

Vertically integrating (3a) using these boundary conditions and (1) gives the continuity equation for vertically averaged biomass

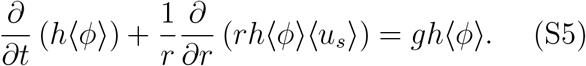

Applying global biomass conservation, the biomass volume 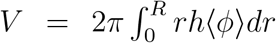 satisfies *∂V/∂t* = *gV*. Expanding this out using the continuity equation (S5), (1) and the boundary conditions in (S4) gives

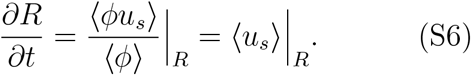

Modelling the upper sheet as a thin elastic beam, the pressure difference across the sheet is

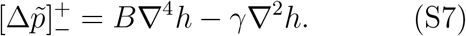

Balancing normal stress at the interface between the biofilm and the sheet yields

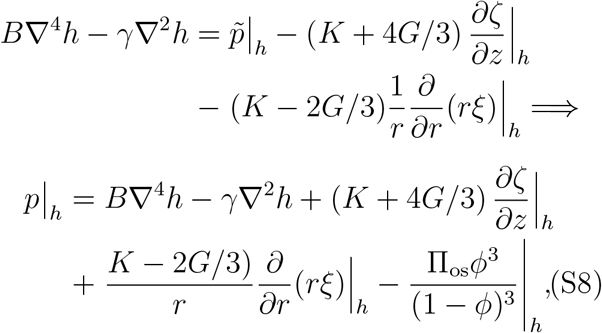

where Π_os_ = *k*_*B*_*T β*^3^*/*3*ν*_0_.

#### Dimensionless shallow-layer scalings

We scale radial and vertical lengths with the initial radius *R*_0_ = *R*(*t* = 0) and height *H*_0_ = *h*(*r* = 0, *t* = 0) of the biofilm respectively i.e {*r, R*} ∼ *R*_0_ and {*z, h*} ∼ *H*_0_. Since the characteristic time scale for the system is that for biofilm growth, we scale *t* ∼ 1*/g*. Utilizing 3) and (S6), we find {*u*_*f*_, *u*_*s*_} ∼ *U*_*s*_ = *gR*_0_ and {*w*_*f*_, *w*_*s*_} ∼ *gH*_0_. Since ***u***_***s***_ is defined as the material derivative of ***ξ***, we have *ξ* ∼ *R*_0_ and *ζ* ∼ *H*_0_. By definition *κ* ∼ *κ*_0_. Finally, a leading order contribution to the pressure comes from the vertical confinement, i.e. (S8) implies 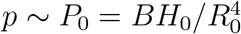. We denote the dimensionless form of a function *f* by *f* ^*^ and set for clarity

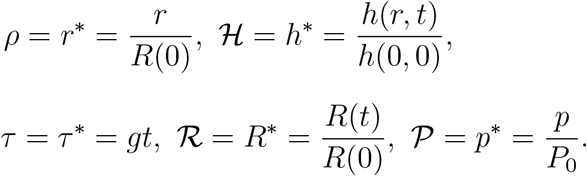

#### Inner Governing Equations

Using these scalings and setting *ϵ* = *H*_0_*/R*_0_, the system of equations (3)-(9) becomes

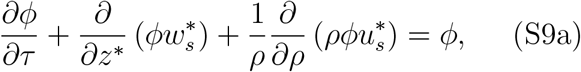

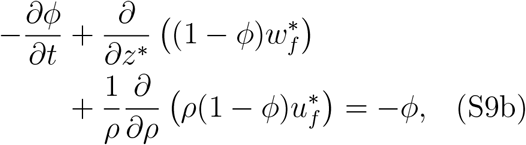

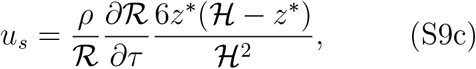

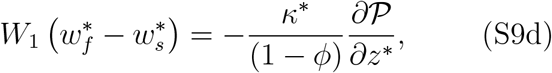

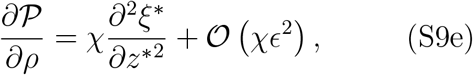

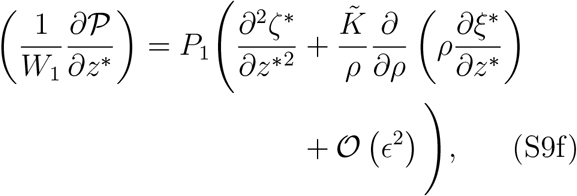

together with the continuity equation for vertically averaged biomass

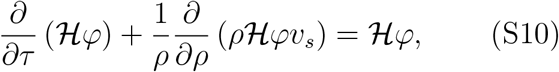

and corresponding vertical boundary conditions

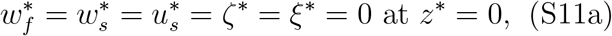

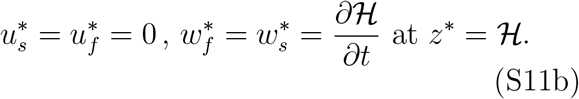

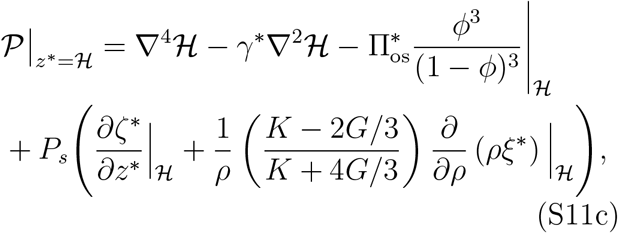

where 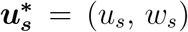 can be expressed as the material derivative of ***ξ***^*^ = (*ξ, ζ*) using

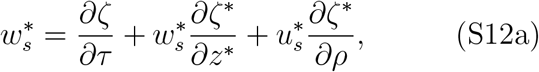

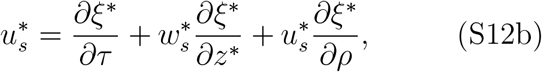

while 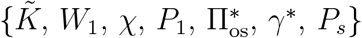 are non-dimensional constants that satisfy

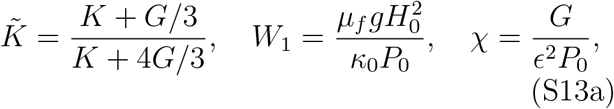

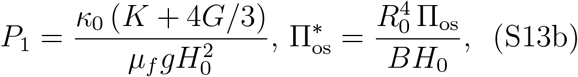

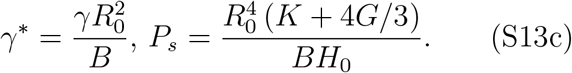

Here, {*W*_1_, *χ, P*_1_} are dimensionless measures of the ability of flow to generate a vertical pressure gradient and the relative strength of the pressure gradients compared to elastic stresses in the horizontal and vertical respectively. *γ*^*^ and 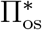 are the non-dimensional surface tension and osmotic pressure scaling groups. Finally, *P*_*s*_ measures the relative strength of the elastic stresses from the biofilm and the PDMS sheet at the upper interface.

#### Order of Magnitude Estimates for Parameters

In a typical experiment, the biofilm initially has height *H*_0_ ∼ 10^−5^m and radius *R*_0_ ∼ 10^−4^m. We assume that the dynamic viscosity of the nutrient rich liquid phase can be approximated by that of water, *μ*_*f*_ ∼ 7.98 × 10^−4^Pa s. From Seminara *et al*., an order of magnitude estimate for the biofilm growth rate *g* is *g*^−1^ ∼ 2.3 h [S1]. Furthermore, the characteristic biofilm permeability scale 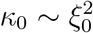 where the biofilm mesh length scale *ξ*_∞_ ∼ 50 nm i.e. *κ*_0_ ∼ 2.5 × 10^−15^m^2^

Picioreanu *et al*. estimated the mechanical properties of a range of different biofilms cultivated from activated sludge supernatant using optical coherence tomography, obtaining an effective Poisson ratio *ν*_*b*_ = 0.4 and Young’s modulus in the range 70 − 700 Pa [S7]. Assuming isotropy, we can thus estimate *K* and *G* as being in the range *K* = *E*_*b*_*/*3(1 − 2*ν*_*b*_) ∼ 117 − 1170 Pa and *G* = *E*_*b*_*/*2(1 + 2*ν*_*b*_) ∼ 19.4 − 194 Pa respectively.

The PDMS sheet has thickness *d* ∼ 10^−4^m, Poisson’s ratio *ν* ∼ 0.5 and Young’s modulus *E* ∼ 1.9 × 10^6^ Pa (a value of 55 measured using a type A durometer). The matrix solid fraction *β* and the volume occupied by one monomer of extracellular matrix varies considerably, depending on a range of factors such as the species of bacteria and the nutrient concentration. Aiming to show that the osmotic pressure contribution can be neglected, we consider uppper and lower bounds for *β* and *ν*_0_ respectively i.e. *β* = 𝒪(1) and *ν*_0_ ∼ 10^−24^m^3^. Finally, we estimate the surface tension between the biofilm and the sheet using that between water and PDMS (*γ* ∼ 4 × 10^−2^Nm^−2^) [S8].

Hence, estimating values for the non-dimensional parameters 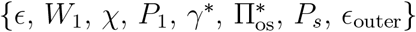 gives

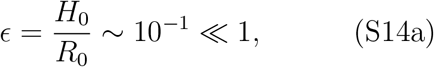

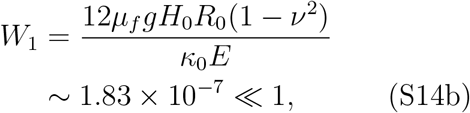

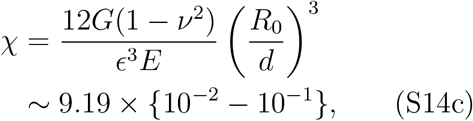

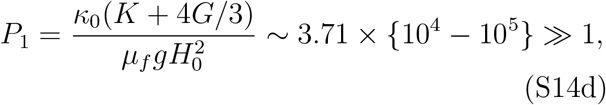

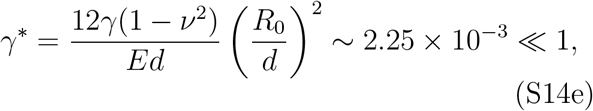

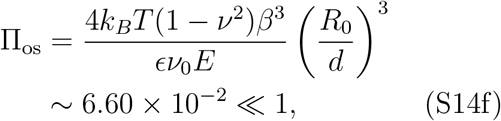

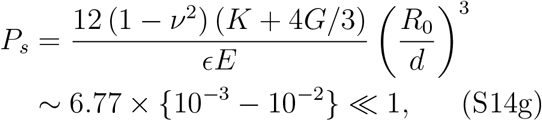

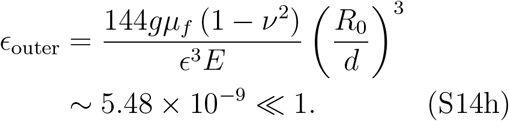

#### Stiff Elastic Confinement

Hence, under experimental conditions, we see that the upper elastic sheet is sufficiently stiff that 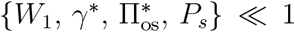 i.e. the dominant contribution to the pressure arises from the upper confinement. *P*_1_ ≫ 1 means that elastic stresses dominate the vertical pressure gradient. In general, *χ* =𝒪(1). Hence, the systems of governing equations given in (*S*9) − (*S*11) reduces to

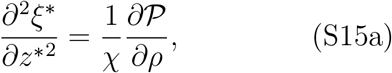

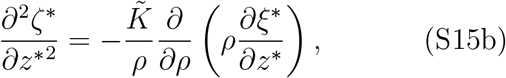

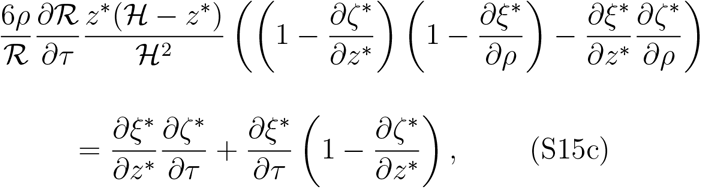

with corresponding boundary conditions

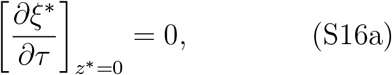

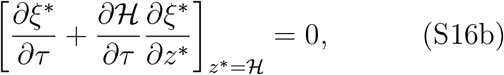

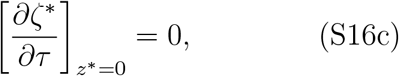

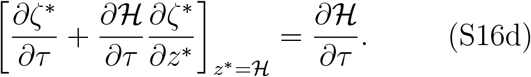

where

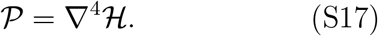

Since 𝒫 is independent of *z*^*^, integrating (S15a) twice with respect to *z*^*^ yields the functional form for *ξ*^*^

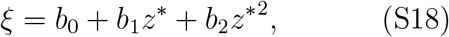

where *b*_*i*_ ={*b*_*i*_(*ρ, τ*) : *i* ∈ [0, 1, 2]} are independent of *z*^*^ and *b*_2_ satisfies

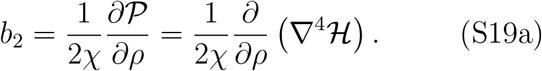

(S16a) and (S16b) simplify respectively to give

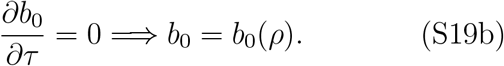

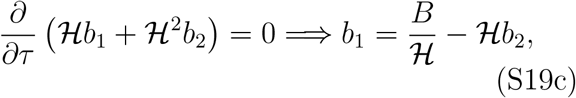

where *B* = *B*(*ρ*) is independent of *τ* and *z*^*^. Hence, (S15b) becomes

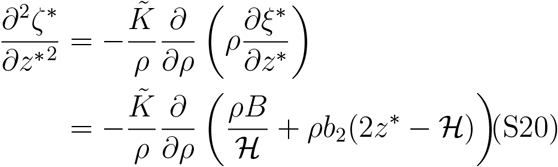

Integrating this twice with respect to *z*^*^ yields the functional form for *ζ*^*^,

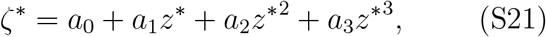

where {*a*_*i*_ = *a*_*i*_(*ρ, τ*) : *i* ∈ [0, 1, 2, 3]} are independent of *z*^*^ and *a*_2_ and *a*_3_ satisfy

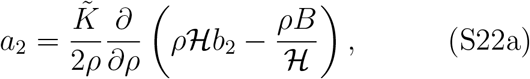

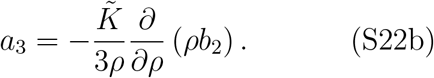

(S16c) and (S16d) simplify respectively to give

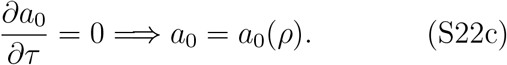

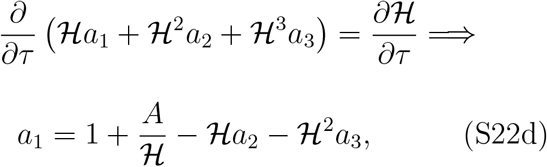

where *A* = *A*(*ρ*) is independent of *τ* and *z*^*^. Substituting (S18) and (S21) into (S15c) and equating the various powers of *z*^*^ gives the following set of six coupled equations for the variables *a*_*j*_ and *b*_*k*_ where *j* ∈ [0, 1, 2, 3] and *k* ∈ [0, 1, 2]

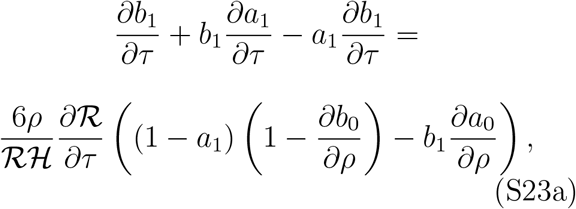

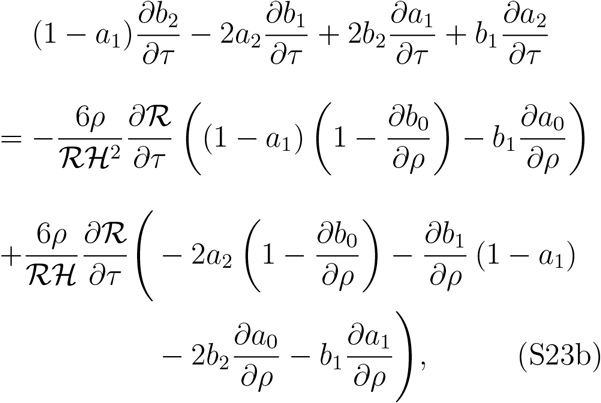

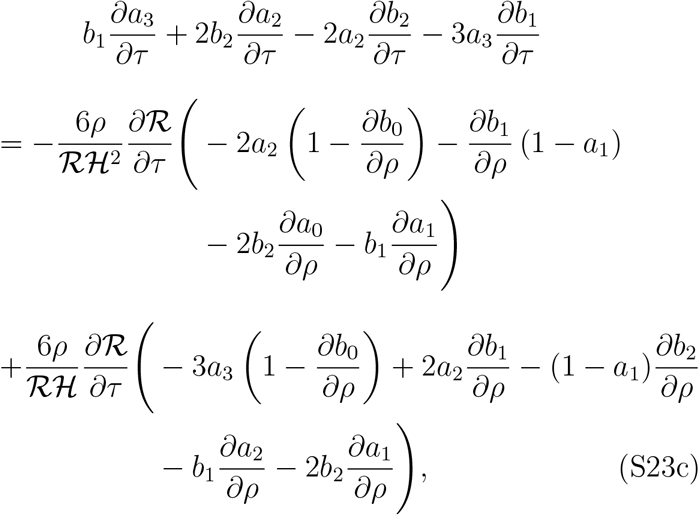

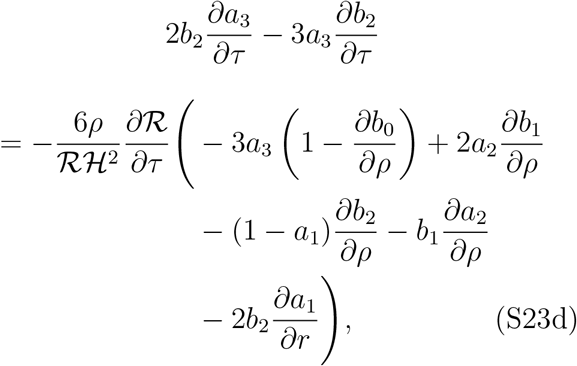

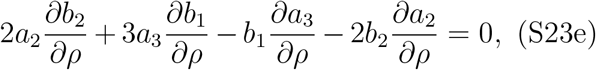

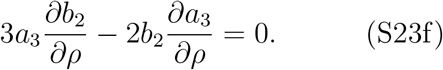

In particular, equating co-efficients of 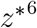 gives

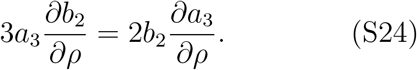

We then have three possible cases:

1. Mode zero, *b*_2_ = 0 =⇒ *a*_3_ = 0,
2. Mode one, *b*_2_ ≠ 0 but *a*_3_ = 0,
3. Mode two, *b*_2_ ≠ 0 and *a*_3_ ≠ 0.

#### Mode zero

When *b*_2_ = 0, ℋ satisfies the differential equation

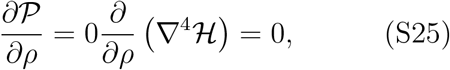

which has the general solution

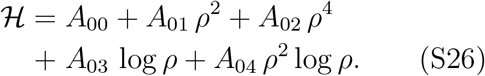

Note that this mode is dominant in the limit *χ* ≪ 1.

#### Mode one

When *a*_3_ = 0, ℋ satisfies the differential equation

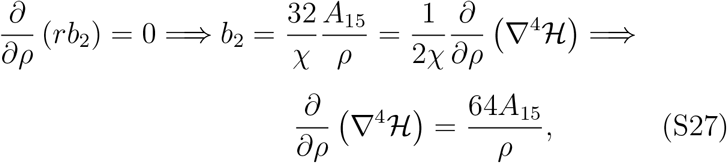

which has the general solution

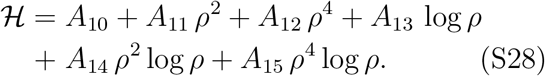

#### Mode two

When both *b*_2_ and *a*_3_ ≠ 0, *b*_2_ satisfies the differential equation

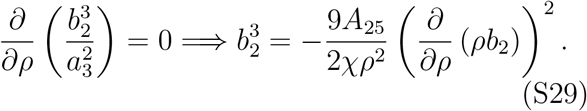

Employing the substitution 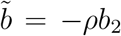, this differential equation becomes separable and can be integrated to give

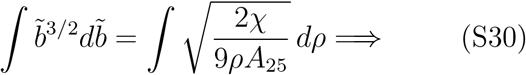

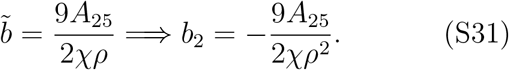

Hence, ℋ satisfies the differential equation

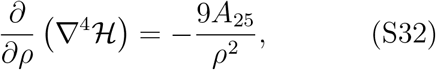

which has the general solution

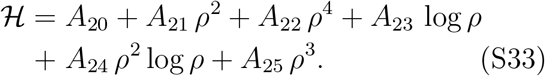

#### Horizontal Boundary Conditions

Define the inverse function of ℛ(*τ*), *τ*_1_(*ρ*), as satisfying

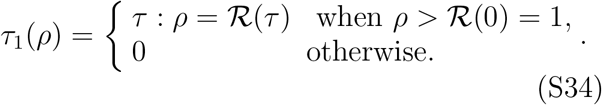

Henc e, constraining the pressure of the solid phase 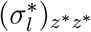 to be constant at the biofilm interface yields

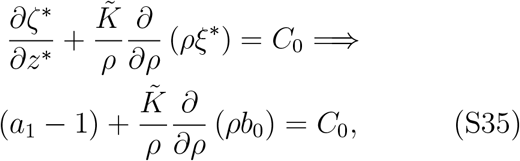

at *τ* = *τ*_1_(*ρ*), where *C*_0_ is a constant which is set from the initial pressure difference at *τ* = 0 across the edge of the biofilm. From symmetry, ℋ and 𝒫 are even in *ρ* at *ρ* = 0, i.e.

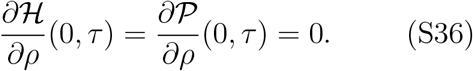

We assume that the biofilm grows uniformly at the interface, namely the vertically averaged biomass volume fraction *φ* Satisfies

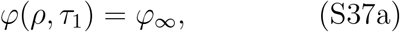

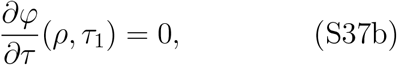

where *φ*_∞_ is a constant.

#### Outer Governing equations

When *ρ > ℛ*, we have a lubrication flow of a single phase Newtonian fluid with viscosity *μ*_*f*_. Hence, the vertically averaged fluid velocity ⟨*u*^*^⟩ satisfies

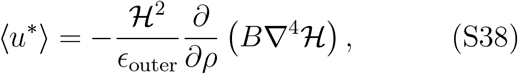

leading to the continuity equation

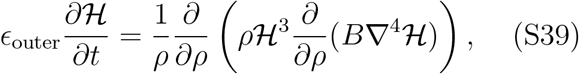

where the nondimensional constant *E*_outer_ satisfies

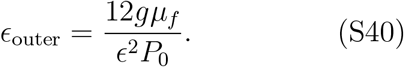

From above, under experimental conditions, *E*_outer_ ≪ 1. Hence, for 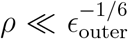 and {*τ*, ℋ} = 𝒪(1), (S39) becomes

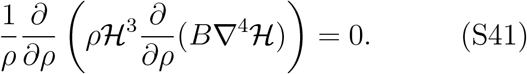

#### Outer Boundary Conditions

In practice, the PDMS sheet has a finite radial extent at *ρ* = ℛ_outer_ = *R*_outer_*/R*(0) where 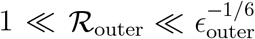. There are two possible kinds of boundary conditions that could be imposed here. If the sheet is clamped, namely fixed height and zero first derivative of height, we have

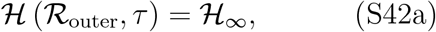

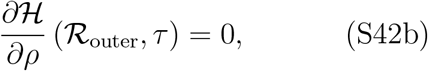

where ℋ_∞_ = *h*_∞_*/H*_0_ is a constant. Alternatively, if the sheet is not clamped, we impose free-beam conditions at *ρ* = ℛ_outer_, namely

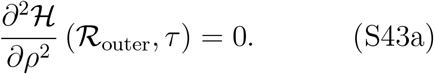

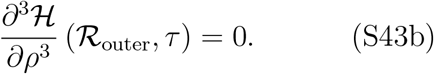

#### Interface Matching Conditions

A fluid flux balance at the biofilm edge yields

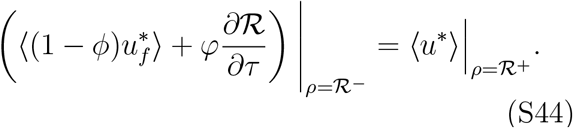

At leading order in *ϵ*_outer_, this simplifies to

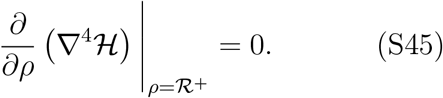

This requires that across *ρ* = ℛ, fourth and lower derivatives of ℋ are continuous.

### SIMPLIFICATION FROM A TWO TO A ONE PHASE SYSTEM

In the above, we have written down a set of governing equations for the full two-phase system, considering both the inner and the outer regions, with corresponding boundary conditions at *ρ* = 0, *ℛ* and ℛ_outer_. Working in the limit that ℛ_outer_ ≫ 1, here we simplify our framework to just considering a single phase system, namely the inner region, together with boundary conditions at *ρ* = 0 and ℛ. Utilising the general form of the solution for ℋ in the outer region, we achieve this by re-writing the far field boundary conditions at *ρ* =ℛ_outer_ (expressed in terms of derivatives of ℋ at *ρ* =ℛ_outer_) in terms of derivatives of ℋ at *ρ* = ℛ, noting that these derivatives are continuous across the biofilm interface.

#### Matching Machinery

Integrating (S41), using the boundary condition given in (S45), the general solution for in the outer region is

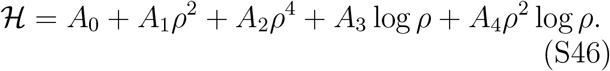

Defining vectors containing the constants of integration, the derivatives of ℋ at the interface and the derivatives of ℋ at the radial extent of the sheet, ***A***_outer_, ℋ_interface_ and ℋ_outer_ respectively, as satisfying

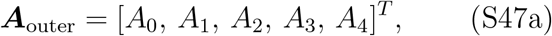

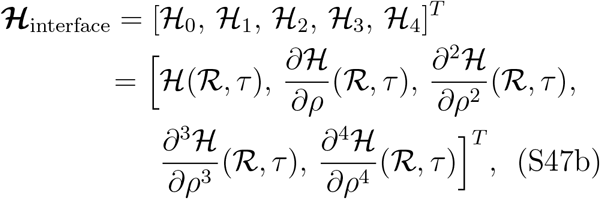

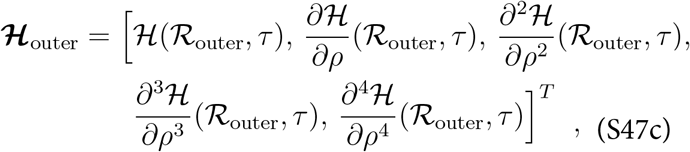

we can express ℋ_outer_ in terms of ℋ_interface_ using (S46):

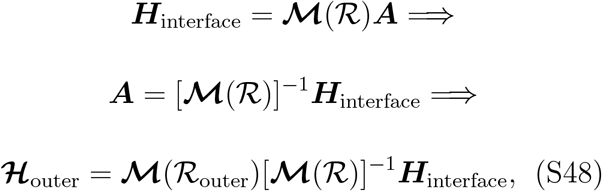

where

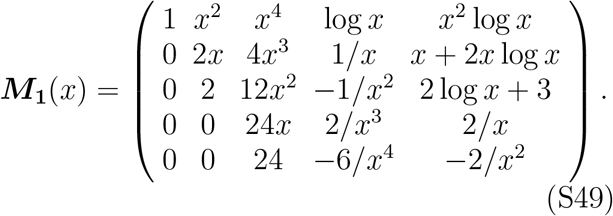

#### Re-writing the Far-field Clamped Boundary Conditions

Using (S48), the boundary conditions at *ρ* = ℛ_*outer*_ given in (S42) can be written in the form

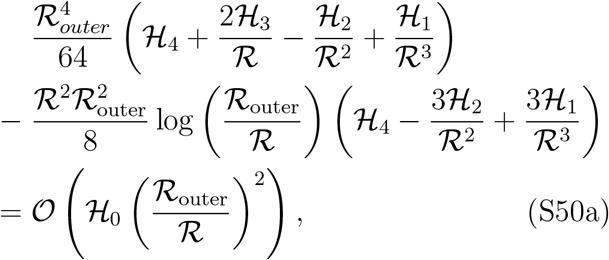

and

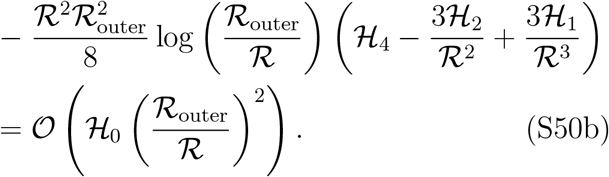

These two conditions rearrange to give

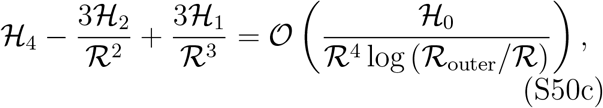

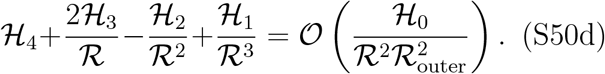

Moving back to tensorial notation, we see that to leading order in ℛ_outer_ the far field boundary conditions can be rewritten as the zero pressure condition

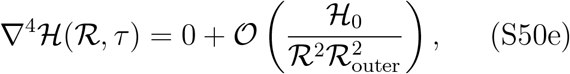

together with the force free condition

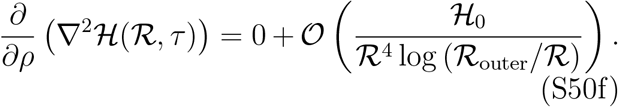

#### Re-writing the Far-field Free Beam Boundary Conditions

In the same way, using (S48), the boundary conditions at *ρ* =ℛ_*outer*_ given in (S43) can be written in the form

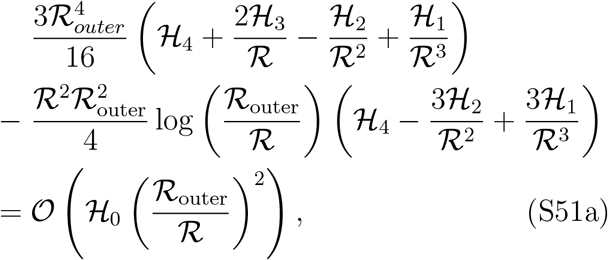

and

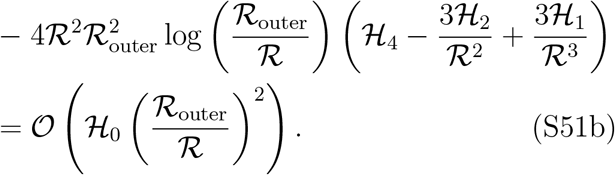

These two conditions rearrange to give

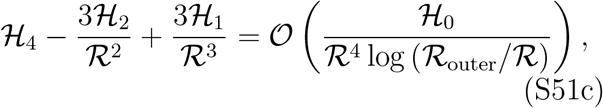

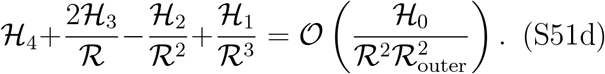

Moving back to tensorial notation, we see that to leading order in ℛ_outer_ the far field boundary conditions can be rewritten as the zero pressure condition

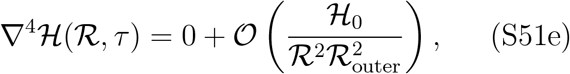

together with the force free condition

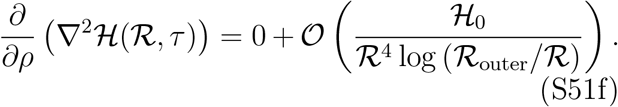

Noting that (S50e), (S50f) and (S51e), (S51f) are identical, we see that both set of boundary conditions at *ρ* = ℛ_outer_, when rewritten in terms of derivatives of ℋ at *ρ* = ℛ, give at leading order in ℛ_outer_ the same conditions for ℋ.

### MODE ZERO SIMILARITY SOLUTION

To make further analytic progress, we look for a similarity solution in the mode zero case i.e.

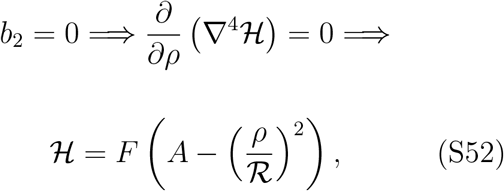

where we have utilised the horizontal boundary conditions for ℋ, *A* is a constant and *F* = *F* (*τ*) is independent of *ρ*. Evaluating (S10) at *ρ* = ℛ then gives

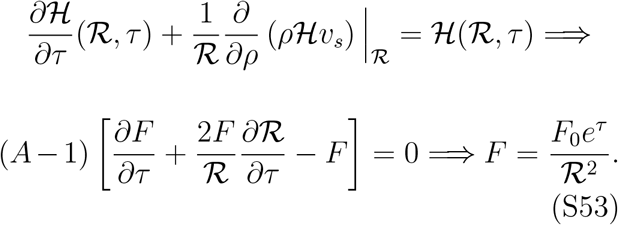

Applying the initial condition ℋ_0_(*ρ* = 0, *τ* = 0) = 1 and defining the incline ratio *m*_0_, a mea-sure of the initial flatness of the biofilm, as sat-isfying

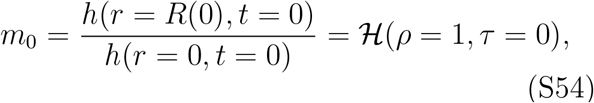

we find *F*_0_ = 1 − *m*_0_ and *A* = 1*/*(1 − *m*_0_) i.e.

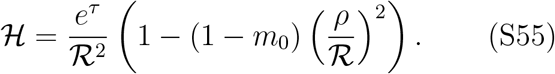

Defining *f* = *f* (*ρ, τ*) = ℋ*φe*^*τ*^, the continuity equation (S10) becomes

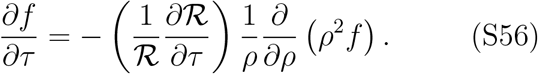

Looking for a similarity solution of the form *f* = *f*_1_(ℛ)*f*_2_(*η*) where *η* = *ρ/ ℛ*, this simplifies to give

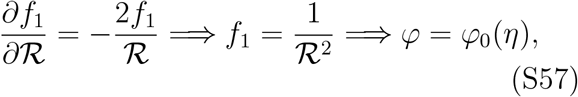

Here *φ*_0_(*ρ*) = *φ*(*ρ, τ* = 0) is set from the initial conditions (*φ*_0_ must satisfy the properties *φ* = *ϕ*_∞_ and *∂φ/∂ρ* = 0 at *τ* = 0).

Finally, we seek an analytical solution for ***ξ***^*^ = (*ξ*^*^, *ζ*^*^) with minimal *z*^*^ dependence. Setting *a*_2_ = 0, (S18) and (S21) simplify to become

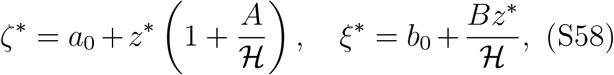

where {*a*_0_, *b*_0_, *A, B*} are all independent of *τ*. (S15a) is automatically satisfied. (S15b) and (S15c) reduce to

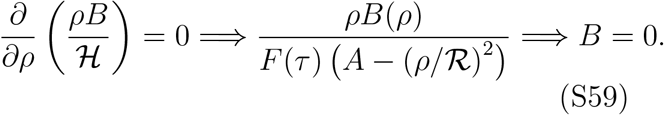

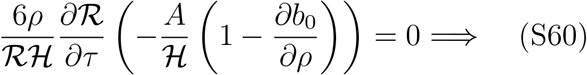

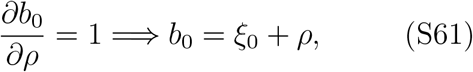

where *ξ*_0_ is a constant set from the initial conditions. Finally, we set for simplicity *A* to a constant *ζ*_0_. Hence, the stress boundary condition (111) simplifies to become

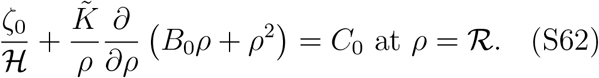

Applying the initial condition ℛ(0) = 1, we recover the cubic equation which describes the evolution of ℛ

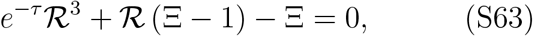

Here, the non-dimensional evolution constant Ξ is

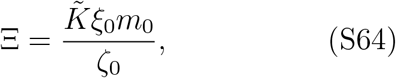

and is thus is determined from the initial conditions as the product of the incline ratio and a ratio between horizontal and vertical stresses.

Now, Cardano’s formula for depressed cubic equations states that for the equation

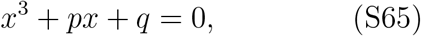

where *p* and *q* are real, if Λ(*p, q*) = 4*p*^3^+27*q*^2^ *>* 0 then the equation has the single real root

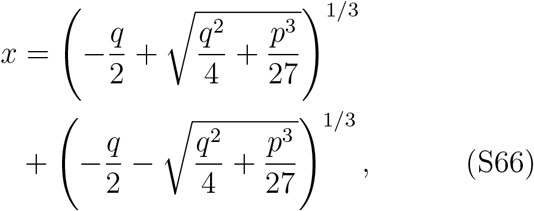

with the other two roots being complex conjugates. If Λ *<* 0 there are three real roots but they can not be represented by an algebraic expression involving only real numbers. This was called by Cardano the *casus irreducibilis* (Latin for ‘the irreducible case’).

For (S63), we have

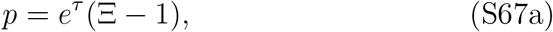

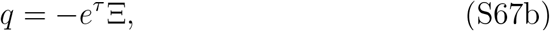

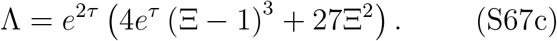

Hence, Λ *<* 0 when Ξ *<* 1 and *τ* satisfies

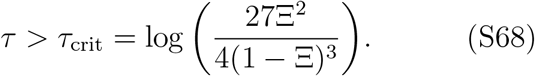

Thus, for general Ξ and *τ*, (S63) does not admit an analytical solution. Instead, this cubic equation is solved numerically using the MATLAB inbuilt function fzero [S6]. Since cubic equations have up to three real roots, we select the correct root by locating the root that is closest to the value for ℛ found at the previous time step, noting that by definition ℛ (*τ* = 0) = 1.

### NEWTONIAN MODEL

Here, for comparison, we analyze the corresponding mathematical model in which, as in Seminara et. al. [S1], the intrinsic elasticity of the biofilm extracellular matrix is neglected. In this case, a solution with power law growth tending to a maximum finite biofilm radius is not supported, demonstrating that matrix elasticity is essential to capture the behavior we have observed experimentally.

#### Dimensionless shallow-layer scalings

In the same way as for the poroelastic model, we scale radial and vertical lengths with the initial radius *R*_0_ = *R*(*t* = 0) and height *H*_0_ = *h*(*r* = 0, *t* = 0) of the biofilm, respectively, permeability with the characteristic permeability scale *κ*_0_, pressure with the vertical confinement pressure scale and time with thath for biofilm growth i.e. 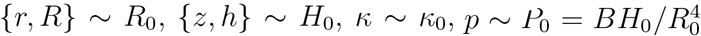 and *t* ∼ 1*/g*. Hence, we find {*u*_*f*_, *u*_*s*_} ∼ *gR*_0_ and {*w*_*f*_, *w*_*s*_} ∼ *gH*_0_. Similarly, we denote the dimensionless form of a function *f* by *f* ^*^ and set for clarity

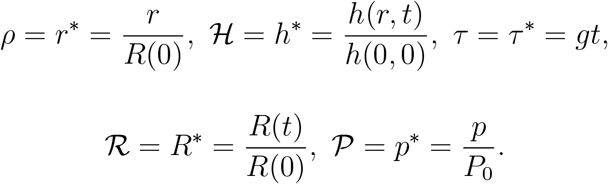

As above, we assume that the biomass volume fraction *ϕ* is independent of *z*^*^,

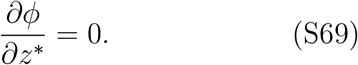

In nondimensional form, the governing equations for this system become

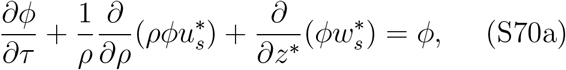

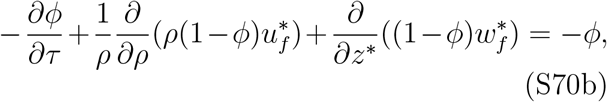

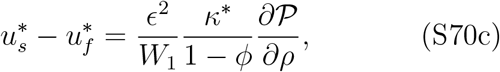

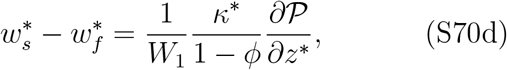

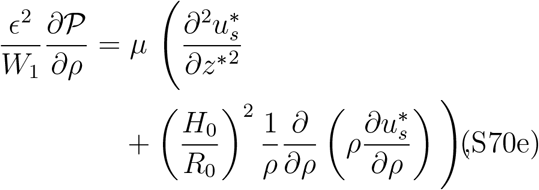

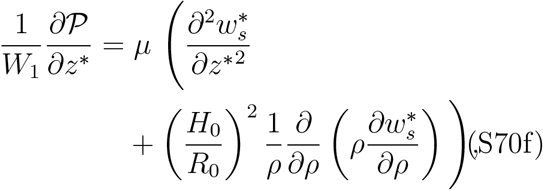

where the non-dimensional constants {*μ, W*_1_} satisfy

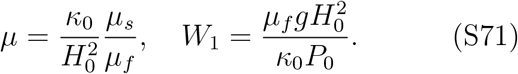

Here *W*_1_, defined in (S13), is a dimensionless measure of the ability of flow to generate a vertical pressure gradient, *μ* is the non-dimensional biofilm viscosity scaling group and *μ*_*s*_ is the dimensional Newtonian viscosity of the biofilm solid phase. Utilising the typical experimental values for the scalings together with *μ*_*s*_ ∼ 10^2^ Pa s we see that {*W*_1_, *ϵ*^−2^*W*_1_} ≪ 1 while *μ* ≈ 2.8 = 𝒪(1).

#### Vertical boundary conditions

As before, imposing no-slip boundary conditions at both the lower and upper boundaries gives

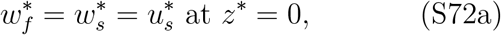

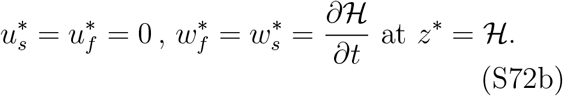

Balancing normal stress at the biofilm sheet interface gives

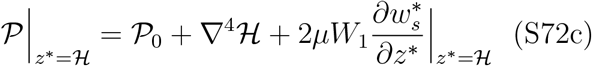

where 𝒫_0_ is a constant reference pressure. Working at leading order in *W*_1_, combining (S70d) and (S72c) gives

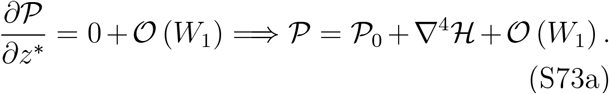

Hence, applying (S70c) gives the differential equation for ℋ

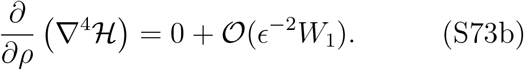

Similarly, combining ((S70d) and (S70f)) and ((S70d) and (S70f)) yields respectively

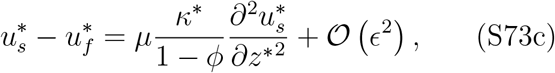

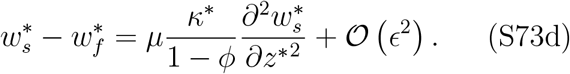

Finally, integrating (S70e) using the boundary conditions given in (S72a) and (S72b) gives

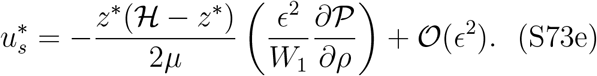

#### Vertically averaged governing equations

We denote vertically averaged quantities by triangular brackets, namely for an arbitrary function *f* we define 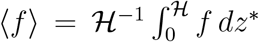, and for clarity set

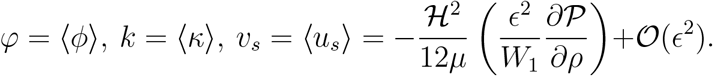

Integrating (S70a) in the *z*^*^ direction yields

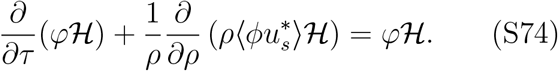

Similarly, integrating (S70a)+(S70b) in the *z*^*^ direction gives the continuity equation

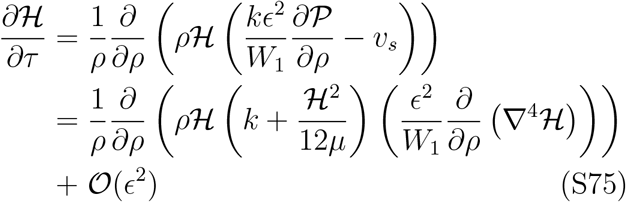

#### Vertically averaged boundary conditions

As in the poroelastic model, we have the boundary conditions

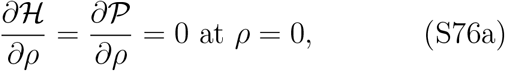

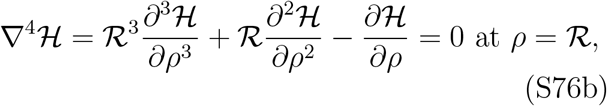

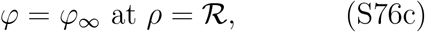

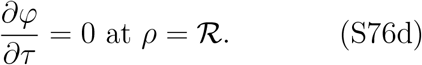

Similarly, polymer volume conservation yields the evolution condition

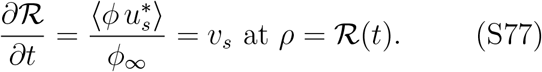

As above, (S73b) together with the boundary conditions in (S76b) admits the similarity solution

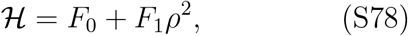

where *F*_0_ = *F*_0_(*τ*) and *F*_1_ = *F*_1_(*τ*) are independent of *ρ*. Integrating (S75) with respect to *ρ* then gives

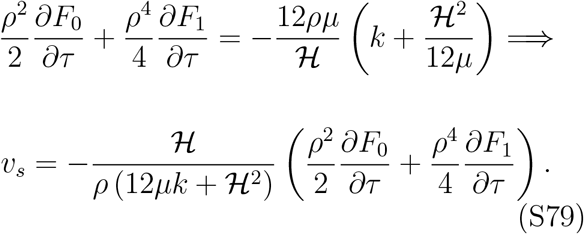

Here, we have used (S76a) to set the integration constant to 0. In general, one can not make further analytic progress.

#### Finite radius solution

Experimentally, we see that the radius of the biofilm tends to a finite value i.e. the system supports a biofilm with constant radius ℛ= ℛ_∞_. In this case, (S77) simplifies to give

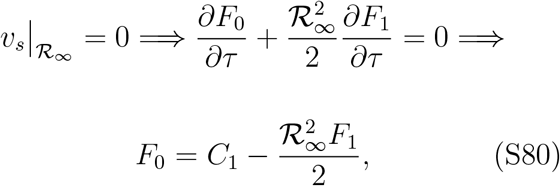

where *C*_1_ is a constant. Since ℋ *>* 0 ∀*ρ* ∈ [0, *ℛ*_∞_], evaluating ℋ at *ρ* = 0 and *ρ* = ℛ_∞_ gives

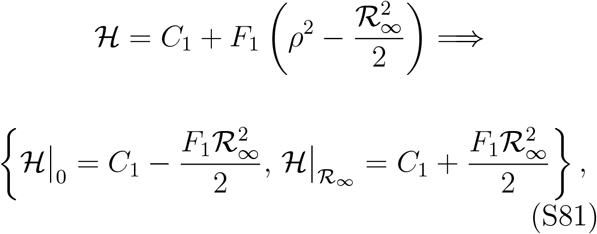

namely *C*_1_ is positive with the lower bound 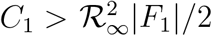. Similarly, differentiating (S79) with respect to *ρ* at *ρ* = ℛ_∞_ gives

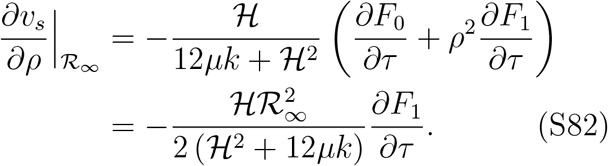

Hence, evaluating (S74) at *ρ* =ℛ, utilising the boundary conditions given above gives

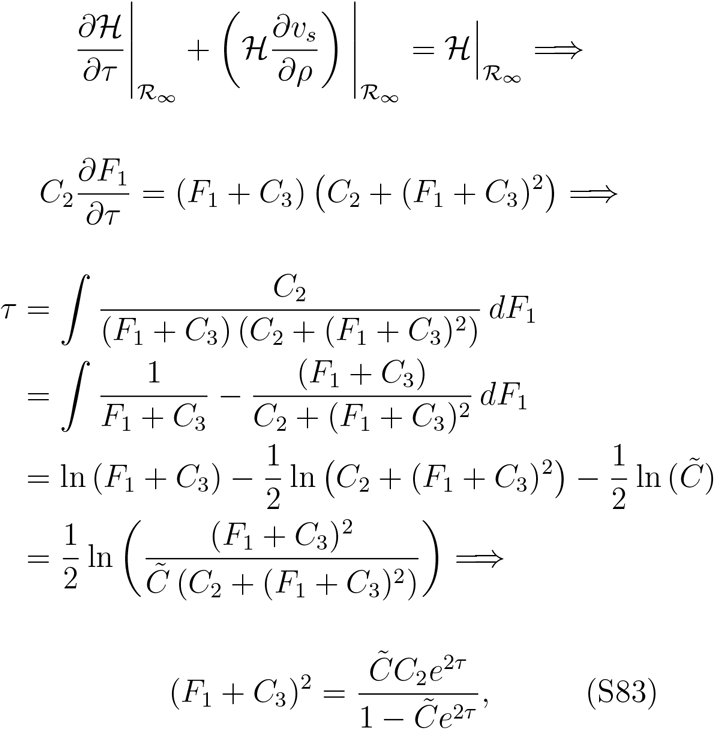

where 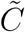 is a constant of integration and the positive constants *C*_2_ and *C*_3_ satisfy

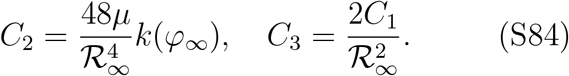

Since the right hand side of (S83) is non-negative for all 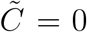 and thus *F*_1_ = −*C*_3_. However, this then gives ℋ = 0 at *ρ* = ℛ_∞_ which is a contradiction. Hence, the Newtonian model does not support a constant radius solution and thus does not agree with experiments.

## Notes

### Competing Interest Statement

The authors have declared no competing interest.

## References

[1] T. Bjarnsholt, Introduction to biofilms, in Biofilm Infections, T. Bjarnsholt, P.ø. Jensen, C. Moser and N. Høiby, eds. (Springer, New York, 2011), pp. 1-9.

[2] R. Hooke, Micrographia, or, Some physiological descriptions of minute bodies made by magnifying glasses :with observations and inquiries thereupon, London : Printed by J. Martyn and J. Allestry, printers to the Royal Society (1665)

[3] P. G. Saraf, A. T. K. Cockett, Marcello Malpighi — A Tribute, Urology 23, 619–623 (1984)

[4] J. Wimpenny, W. Manz, U. Szewzyk, Heterogeneity in biofilms, FEMS Microbiol. Rev. 24, 661 (2000).

[5] A. van Leeuwenhoek, An abstract of a letter from Mr. Anthony Leewenhoeck at Delft, dated Sep. 17. 1683. Containing some microscopical observations, about animals in the scurf of the teeth, the substance call’d worms in the nose, the cuticula consisting of scales, Phil. Trans. R. Soc. 14, 568 (1684).

[6] J. W. Costerton, Z. Lewandowski, D. Debeer, D. Caldwell, D. Korber, G. James, Biofilms, the Customized Microniche, J Bacteriol. 176, 2137 (1994).

[7] G. O’Toole, H. B. Kaplan, R. Kolter, Biofilm formation as microbial development, Annu. Rev. Microbiol. 54, 49–79 (2000)

[8] D. López, H. Vlamakis, and R. Kolter, Biofilms, Cold Spring Harb. Perspect. Biol. 2, a000398 (2010).

[9] A. Seminara, T.E. Angelini. J.N. Wilking, H. Vlamakis, S. Ebrahim, R. Kolter, D.A. Weitz, and M. P. Brenner, Osmotic spreading of Bacillus subtilis biofilms driven by an extracellular matrix, Proc. Natl. Acad. Sci. USA 109, 1116 (2012).

[10] A. Tam, E.F. Green, S. Balasuriya, E.L. Tek, J.M. Gardner, J.F. Sundstrom, V. Jiranek, and B.J. Binder, A thin-film extensional flow model for biofilm expansion by sliding motility, Proc. R. Soc. A 475, 20190175 (2019).

[11] S. Srinivasan, C.N. Kaplan, and L. Mahadevan, A multiphase theory for spreading microbial swarms and films, eLife 8, e42697 (2019).

[12] R. Martinez-Corral, J. Liu, A. Prindle, G. M. Süel, and J. Garcia-Ojalvo, Metabolic basis of brain-like electrical signalling in bacterial communities, Phil. Trans. R. Soc. B 374, 20180382 (2019).

[13] F. Kempf, R. Mueller, E. Frey, and J. M. Yeomans, Active matter invasion, Soft Matter 15, 7538 (2019).

[14] J. C. Conrad and R. Poling-Skutvik, Confined Flow: Consequences and implications for bacteria and biofilms, Annu. Rev. Chem. Biomol. Eng. 9, 175 (2018).

[15] Z. Khatoon, C.D. McTiernan, E.J. Suuronen, T.-F. Mah, and E. I. Alarcon, Bacterial biofilms formation on implantable devices and approaches to its treatment and prevention, Heliyon 4, 1 (2018).

[16] Y. Oppenheimer-Shaanan, N. Steinberg, and I. Kolodkin-Gal, Small molecules are natural triggers for the disassembly of biofilms, Trends Microbiol. 21, 594 (2013).

[17] O. Ciofu, E. Rojo-Molinero, M.D. Macía, and A. Oliver, Antibiotic treatment of biofilm infections, APMIS 125, 304 (2017).

[18] P. S. Stewart, Mechanisms of antibiotic resistance in bacterial biofilms, Int. J. Med. Microbiol. 292, 107 (2002).

[19] C. R. Arciola, D. Campoccia, and L. Montanaro, Implant infections: adhesion, biofilm formation and immune evasion, Nat. Rev. Microbiol. 16, 397 (2018).

[20] J. Nowakowska, R. Landmann, and N. Khanna, Foreign body infection models to study host-pathogen response and antimicrobial tolerance of bacterial biofilms, Antibiotics 3, 378 (2014).

[21] S.S.L. Peppin, J.A.W. Elliott, and M. G. Worster, Pressure and relative motion in colloidal suspensions, Phys. Fluids 17, 053301 (2005).

[22] H.F. Wang, Theory of Linear Poroelasticity with Applications to Geomechanics and Hydrogeology (Princeton University Press, Princeton, 2001).

[23] D.R. Hewitt, J.A. Neufeld, and N.J. Balmforth, Shallow, gravity-driven flow in a poro-elastic layer, J. Fluid Mech. 778, 335 (2015).

[24] R.E. Gibson, R.L. Schiffman, and S.L. Pu, Plane strain and axially symmetric consolidation of a clay layer on a smooth impervious base, Q. J. Mech. Appl. Maths 23, 505 (1970).

[25] S.I. Barry, G.N. Mercer, and C. Zoppou, Deformation and fluid flow due to a source in a poro-elastitc layer, Appl. Math. Model. 21, 681 (1997).

[26] G.T. Charras, T.J. Mitchison, and L. Mahadevan, Animal cell hydraulics, J. Cell Sci. 122, 3233 (2009).

[27] P.J. Flory, Principles of Polymer Chemistry (Cornell University Press, Ithaca, 1953).

[28] H.F. Winstanley, M. Chapwanya, M.J. McGuinness, and A.C. Fowler, A polymer-solvent model of biofilm growth, Proc. R. Soc. A 467, 1449 (2011).

[29] Note that due to the dead volume occupied by the bacteria cells, the extracellular matrix occupies a volume ϕ_m_ within a total volume of 1−ϕ and thus we expand in terms of ϕ_m_/1−ϕ rather than ϕ_m_.

[30] See Supplemental Material at http://link.aps.org/supplemental/10.1103/XXX for further theoretical details and experimental methods, which includes Refs. p31-36.

[31] J. Liu, A. Prindle, J. Humphries, M. Gabalda-Sagarra, D.D. Lee, S. Ly, J. Garcia-Ojalvo, and G. M. Süel, Metabolic co-dependence gives rise to collective oscillations within biofilms, Nature 523, 550 (2015).

[32] J. Humphries, L. Xiong, J. Liu, A. Prindle, F. Yuan, H.A. Arjes, L. Tsimring, and G.M. Süel, Species-independent attraction to biofilms through electrical signaling, Cell 168, 200 (2017).

[33] J. Schindelin, I. Arganda-Carreras, E. Frise, V. Kaynig, M. Longair, T. Pietzsch, S. Preibisch, C. Rueden, S. Saalfeld, B. Schmid, J.-Y. Tinevez, D.J. White, V. Hartenstein, K. Eliceiri, P. Tomancak, and A. Cardona, Fiji: an open-source platform for biological-image analysis, Nature Methods 9, 676 (2012).

[34] R. P. Brent, Algorithms for Minimization Without Derivatives (Prentice-Hall Inc. 1973).

[35] C. Picioreanu, F. Blauert, H. Horn, and M. Wagner, Determination of mechanical properties of biofilms by modelling the deformation using optical coherence tomography, Water Res. 145, 588–598 (2018).

[36] A.E. Ismail, G.S. Grest, D.R. Heine, and M. J. Stevens, Interfacial Structure and Dynamics of Siloxane Systems: PDMS-Vapor and PDMS-Water, Macromolecules 42, 3186–3194 (2009).

## References

[S1] A. Seminara, T.E. Angelini. J.N. Wilking, H. Vlamakis, S. Ebrahim, R. Kolter, D.A. Weitz, and M. P. Brenner, Osmotic spreading of Bacillus subtilis biofilms driven by an extracellular matrix, Proc. Natl. Acad. Sci. USA 109, 1116 (2012).

[S2] R. Martinez-Corral, J. Liu, A. Prindle, G. M. Süel, and J. Garcia-Ojalvo, Metabolic basis of brain-like electrical signalling in bacterial communities, Phil. Trans. R. Soc. B 374, 20180382 (2019).

[S3] J. Liu, A. Prindle, J. Humphries, M. Gabalda-Sagarra, D.D. Lee, S. Ly, J. Garcia-Ojalvo, and G.M. Süel, Metabolic co-dependence gives rise to collective oscillations within biofilms, Nature 523, 550 (2015).

[S4] J. Humphries, L. Xiong, J. Liu, A. Prindle, F. Yuan, H.A. Arjes, L. Tsimring, and G.M. Süel, Species-independent attraction to biofilms through electrical signaling, Cell 168, 200 (2017).

[S5] J. Schindelin, I. Arganda-Carreras, E. Frise, V. Kaynig, M. Longair, T. Pietzsch, S. Preibisch, C. Rueden, S. Saalfeld, B. Schmid, J.-Y. Tinevez, D.J. White, V. Hartenstein, K. Eliceiri, P. Tomancak, and A. Cardona, Fiji: an open-source platform for biological-image analysis, Nature Methods 9, 676 (2012).

[S6] R. P. Brent, Algorithms for Minimization Without Derivatives (Prentice-Hall Inc. 1973).

[S7] C. Picioreanu, F. Blauert, H. Horn, and M. Wagner, Determination of mechanical properties of biofilms by modelling the deformation using optical coherence tomography, Water Res. 145, 588–598 (2018).

[S8] A.E. Ismail, G.S. Grest, D.R. Heine, and M. J. Stevens, Interfacial Structure and Dynamics of Siloxane Systems: PDMS-Vapor and PDMS-Water, Macromolecules 42, 3186–3194 (2009).

